# Computational multiphase characterization of perfusion trends inside biomimetic reduced-order dense tumors

**DOI:** 10.1101/2022.05.30.494072

**Authors:** Mohammad Mehedi Hasan Akash, Nilotpal Chakraborty, Jiyan Mohammad, Katie Reindl, Saikat Basu

**Affiliations:** Department of Mechanical Engineering, South Dakota State University, Brookings, SD 57007, United States; Department of Biomedical Engineering and Mechanics, Virginia Tech, Blacksburg, VA 24061, United States; Center for Diagnostic and Therapeutic Strategies in Pancreatic Cancer, North Dakota State University, Fargo, ND 58108, United States; Department of Biological Sciences, North Dakota State University, Fargo, ND 58108, United States

**Keywords:** Solid tumor, Multiphase simulation, Plasma perfusion, Computational modeling, Biomimetic analysis

## Abstract

Dense fibrous extracellular constitution of solid tumors exerts high resistance to diffusive transport into it; additionally, the scarcity of blood and lymphatic flows hinders convection. The complexity of fluidic transport mechanisms in such tumor environments still presents open questions with translational end goals. For example, clinical diagnosis and targeted drug delivery platforms for such dense tumors can ideally benefit from a quantitative framework on plasma uptake into the tumor. In this study, we present a computational model for physical parameters that may influence blood percolation and penetration into a simple biomimetic solid tumor geometry. The model implements 3-phase viscous laminar transient simulation to mimic the transport physics inside a tumor-adhering blood vessel and measures the constituent volume fractions of the three considered phases, *viz*. plasma, RBCs (Red Blood Cells, also known as “erythrocytes”), and WBCs (White Blood Cells, also known as “leukocytes”) at three different flow times, while simultaneously recording the plasma pressure and velocity at the entry point to the tumor’s extracellular space. Subsequently, to quantify plasma perfusion within the tumor zone, we have proposed a reduced-order 2D transport model for the tumor entry zone and its extracellular space for three different fenestra diameters: 0.1, 0.3, and 0.5 *μ*m; the simulations were 2-phase viscous laminar transient. The findings support the hypothesis that plasma percolation into the tumor is proportional to the leakiness modulated by the fenestra openings, quantifiable through the opening sizes.

## 1. Introduction

Perfusion mechanics into a dense solid tumor has long been recognized as a critical issue in both clinical and experimental studies (Gullino, PM. 1980, Jain, R. K. 1997, Vaupel, P.and Kallinowski, F. and Okunieff, P. 1990). Owing to constriction of tumor blood vessels and reduced leakiness into the tumor extracellular space from the tumor vasculature, blood perfusion in tumors can be much lower than in normal tissues surrounding them (Stylianopoulos, T. and Jain, R. K. 2013). Solid tumors present an abnormal mass of tissues that does not contain cysts or liquid regions, and area categorized as benign (non-cancerous) or malignant (cancerous) (Gavhane, YN. et al. 2011). These tumors exhibit high resistance to diffusive transport (Jain, R. K. 1996, 1998), and the scarcity of blood and lymphatic flow in the surrounding environment impedes convection. These nuances in flow physics render it difficult for blood to perfuse into the tumor volume (Chim, L. K. and Mikos, A. G. 2018), despite the increased permeability and retention effects frequently observed during nanoparticle delivery to tumor tissues (Maeda 2012). Accurate quantification of the spatiotemporal distribution of particulates in the blood channel and tumor region could be critical for cancer clinical diagnosis (d’Esposito, A. et al. 2018). However, the pathophysiology of diseased tissues interacts chaotically with the blood uptake process (Soltani, M. and Chen, P. 2011), and, on a macroscale, can vary between subjects resulting from the variations in tumor topography (Zhang, Y. and Chen, L. et al. 2013).

Review of the biomedical science literature demonstrates that single-phase computational fluid dynamics (CFD) models, which treat blood as a homogeneous fluid (Attinger, EO. 1964, Womersley, JR. 1957), cannot provide the hemodynamic information necessary to quantify the interactive effects and spatial distribution of suspended particulate matters in blood (Srivastava, VP. 2007). In practice, blood exhibits a non-Newtonian rheology (Merrill, E. W. et al. 1963) owing to the complex strain rates as well as high cell count and particulate nature of RBCs (Red Blood Cells). For concentrated suspensions (Baskurt, O. K. and Meiselman, H. J. 2003), the spatial variation of blood constituents such as plasma, RBCs, and WBCs (White Blood Cells) may vary owing to disturbed flow within the tumor region. RBCs collide with one another in such flows as a result of their relative motion and aggregate to form larger RBC agglomerates. This phenomenon, referred to as RBC aggregation, is just one of the inter-particle phenomena of blood flow that affects the rheological behavior of blood and thus the ambient distribution of particulate matter. As a result, developing a more realistic CFD model is critical to realistically mimicking these types of phenomena.

Our model of blood transport in the tumor vaculature is constructed as a multiphase system composed of two dispersed phases, RBCs and WBCs suspended in plasma. Our model is an extension of Gidaspow, D. 1994’s and Anderson, T. B. and Jackson, R. 1967’s multiphase CFD approach. This method is particularly effective for describing blood flows containing closely spaced RBCs, which have a volume fraction of between 30% and 55% *in vivo* (Jung et al. 2006). In comparison to the single-phase model, the multiphase model includes the volume fraction of each phase as well as mechanisms for momentum exchange.

While perfusion into the tumor has been identified as a significant issue for cancer research, herein we attempt to answer the following hypothesis-driven question: can we spot the impact of tumor leakiness on plasma perfusion into the tumor via our reduced-order modeling approach with a 2D biomimetic tumor domain, through considering different sizes of fenestra openings on the tumor vasculature?

To answer the aforementioned, we have used CFD techniques to model the blood transport process in the generated biomimetic model. This study’s preliminary findings were presented at the American Physical Society’s Division of Fluid Dynamics Annual Meeting 2021 (Akash, M. M. H. et al. 2021).

## 2. Methods

### 2.1. Development of biomimetic model

The extracellular matrix supports the biological tissues mechanically, and fiber alignment (Eekhoff, J. D. and Lake, S. P. 2020) influences cellular behavior. Published computed tomographic reconstructions (see panel (a), Figure 1) and a 2D slice through our subsequent 3D idealization (panel (b), Figure 1) serves as the inspiration for the *in silico* 2D (see panel (a), Figure 2) biomimetic geometry used in this study. The orientation of preliminary 3D model’s internal fiber bundles was determined using panel (a) of Figure 1, which was adapted from Eekhoff, J. D. and Lake, S. P. 2020’s work. The 2D biomimetic idealization was built on the Workbench 2019 R3 New Design Modeler Geometry (DM) (ANSYS Inc., Canonsburg, Pennsylvania). We divided this 2D model into two parts to restrict the entry of bigger particles into the smaller path; e.g., to restrict the entry of RBCs (the diameter (Nithiarasu 2022) is 7 *μ*m) into the fenestra (the maximum diameter in our model is 0.5 *μ*m). The two parts are: (i) the blood vessel with depression signifying the entry location to the tumor extracellular space; and (ii) the tumor extracellular domain with a fenestra opening. The depth of the depression was 2.6 *μ*m, and the diameter was 5.2 *μ*m. The depression depth is chosen to remain greater than the height of the mean fenestra opening (i.e., 0.3 *μ*m) in order to ensure that the entire fenestra is contained within the depression region, which is critical for making the biomimetic model realistic as the fenestra serves as the link between the blood vessel and the tumor region. The diameter of the depression is likewise determined with the assumption that the fenestra is completely fitted and does not become outliered, which is why the diameter stays nearly ten-fold that of the fenestra’s maximum diameter (i.e., 0.5 *μ*m) in the biomimetic model. The zoomed views of the depression in the blood vessel is shown in panels (b)-(c) of Figure 2. Figure 2’s panel (d) depicts the tumor domain with fenestra. We modeled three distinct tumor domain geometries using three different yet realistic fenestra diameters of 0.1 *μ*m (Model 1), 0.3 *μ*m (Model 2), and 0.5 *μ*m (Model 3), with the streamwise fenestra length remaining constant at 0.3 *μ*m in each case. Panels (b)-(e) in Figure 2 show the finer mesh elements, which were generated on ANSYS Workbench 2019 R3 Mesh. Each computational grid in this study has more than 0.02 million unstructured and graded octahedral components, with 0.04 million for blood vessel (see panel (b), Figure 2), and 0.023 million for the tumor geometry (see panel (d), Figure 2).

**Fig. 1:**
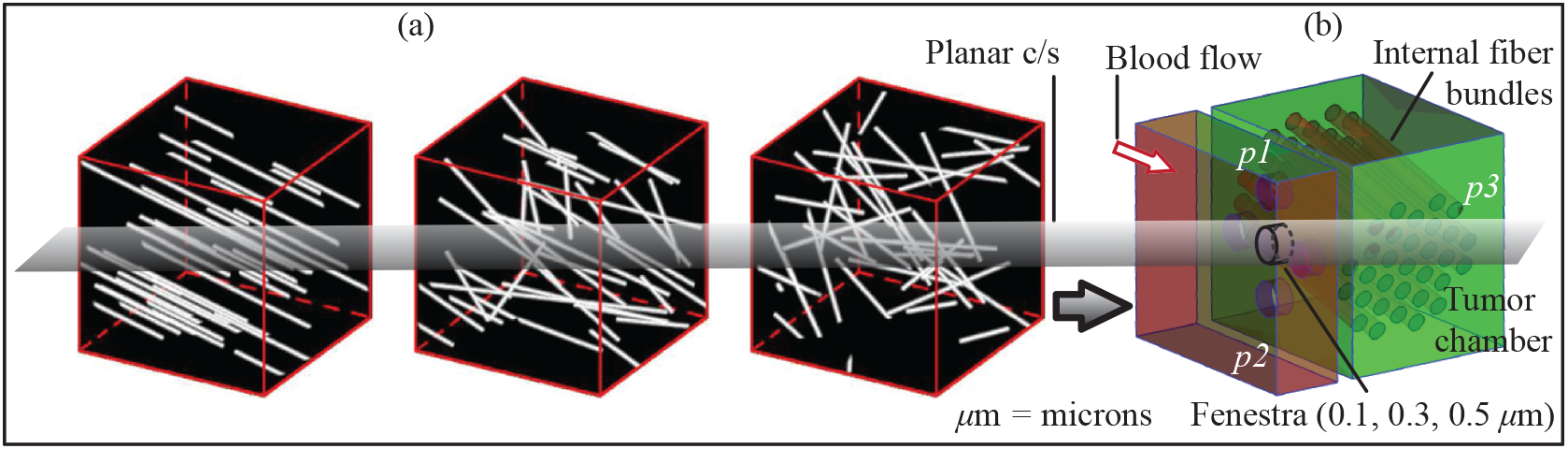
Panel (a) shows the extracellular fiber matrix in scanned tumor domains (source: Eekhoff, J. D. and Lake, S. P. 2020) and panel (b) depicts our preliminary 3D biomimetic concept with random fiber orientation. Permission is granted from Jeremy D. Eekhoff and Spencer P. Lake, 2020, © Royal Society to re-use their work in panel (a).

**Fig. 2:**
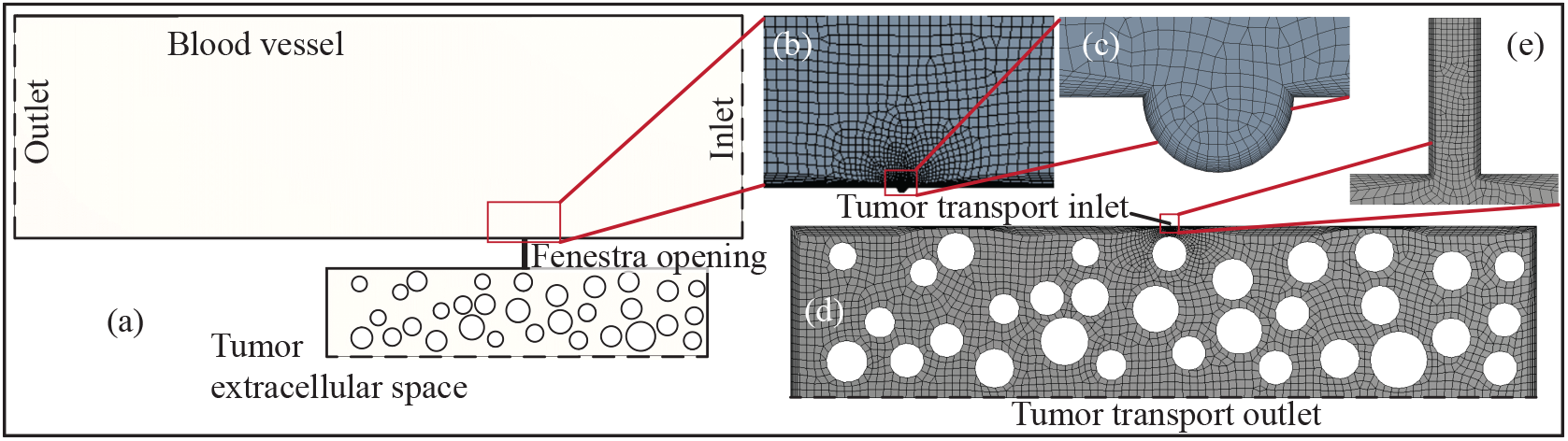
Panel (a) shows the 2D biomimetic model where blood vessel is connected with tumor region by a fenestra. Then we divided panel (a) into two segments to “restrict” the entry of larger particles inside the fenestra which is smaller than the particle sizes. Panels (b) and (d) show those two segments with finer mesh elements. Therein, panel (b) shows the blood vessel with depression; panel (c) zooms in on the artificial depression and panel (d) shows the tumor region with fenestra; panel (e) depicts the fenestra with finer mesh elements. The dashed bottom line in (a) and (d) depict the outlet sink for tracking the intra-tumoral uptake.

### 2.2. Simulation of blood transport inside the blood vessel

The viscous-laminar-transient state flow physics dominates the *in silico* transport of blood inside blood vessels. The system was numerically tracked through a 3D Eulerian multiphase model, wherein we introduced user-defined functions for the shear thinning viscosity of RBCs and the effect of RBC agglomeration using equations from Jung, J. and Hassanein, A. 2008. The effective RBC shear viscosity, *η*_*RBC*_, was calculated from the equation for dimensionless relative blood mixture viscosity, *μ*_*mix*_, in a non-Newtonian shear thinning fluid:

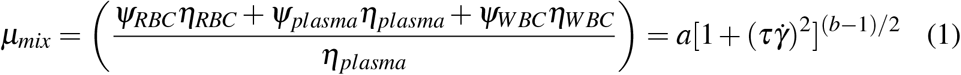

Here 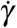 is the shear rate (1/sec). The volume fraction and viscosity are represented by *Ψ* and *η* respectively. The time constant, *τ*, was taken to be 0.110 sec and the two parameters a and b as a function of the hematocrit, H = *Ψ*_*RBC*_, were given by the following polynomial approximations for 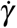 greater than or equal to 6 (Jung, J. et al. 2006).

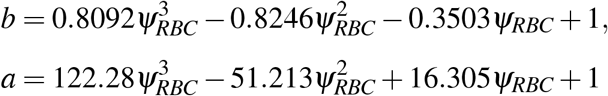

For 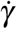 less than 6 at low shear rates, these were as follows:

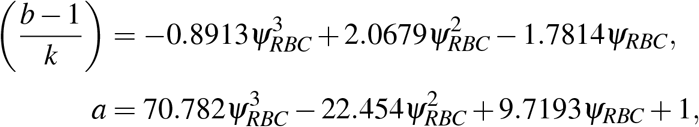

where, 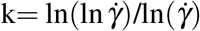

The inlet velocity is pulsatile with a time period of 0.735 sec (see Figure 3). To match it with an actual heart cycle, published experimental data (Cholley, B. P. et al. 1995, Nichols, W. W. 1998, Poppas, A. et al. 1997) is mapped into a Fourier transform with MATLAB curve fitting tool. Both RBCs and WBCs experience interphase drag from plasma, which is a type of fluid-particle interaction. This interaction was described using the interphase momentum exchange coefficients, *α*, and the velocity differences between the continuous phase 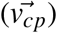 and the dispersed phase 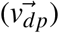.

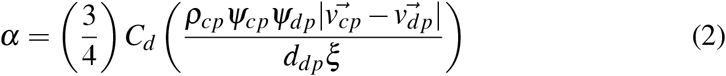

where, the density and volume fraction of the continuous phase are *ρ*_*cp*_ and *Ψ*_*cp*_ respectively, and the volume fraction of the dispersed phase is *Ψ*_*dp*_. The drag coefficient, *C*_*d*_, on a single sphere is related to the Reynolds number, *Re*_*p*_, as per the well known Schiller and Naumann drag model:

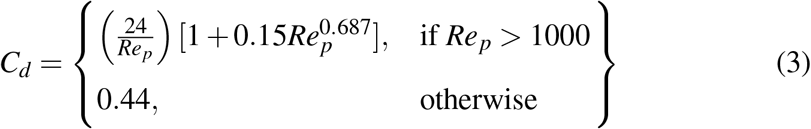

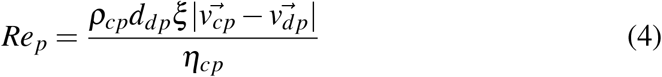

where *d*_*dp*_ is the diameter of blood cells in the dispersed phase, *η*_*cp*_ is the viscosity of the continuous phase, and *ξ* is the dynamic shape factor. More drag is experienced by an agglomerate non-spherical particles in a fluid (Iimura, K. and Higashitani, K. 2005). The dynamic shape factor of RBCs was used to describe the effect of RBC agglomeration as:

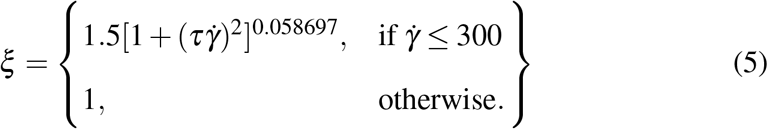

**Fig. 3:**
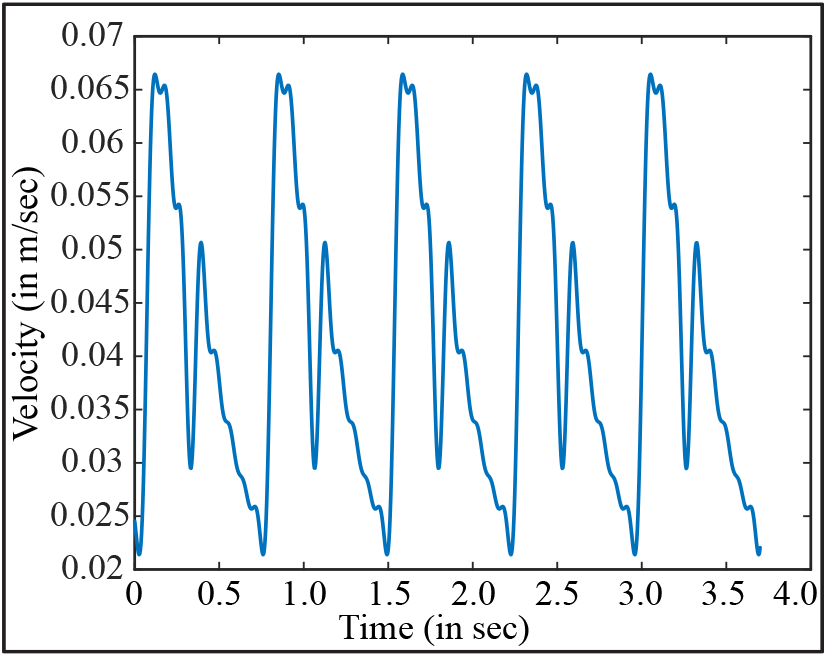
Inlet velocity corresponding to an actual heart cycle. The periodic profile was generated by mapping a Fourier transform from experimental data (Cholley, B. P. et al. 1995, Nichols, W. W. 1998, Poppas, A. et al. 1997)

Similarly the shape factor for WBCs is assumed to be 1 always as they do not undergo agglomeration like RBCs.

Two external forces, *viz*. virtual mass and lift force, were neglected because they are minimal in comparison to the drag force. The numerical solution method makes use of the implicit, unstructured mesh of finite volume elements. The Eulerian multiphase model for tracking transport inside the tumor-adhering blood vessel incorporated the following three phases: plasma, RBCs, and WBCs. Therein, plasma is classified as the primary phase, while RBCs and WBCs are classified as the secondary phases.

The simulations employed a segregate solver, along with SIMPLEC pressurevelocity coupling and first order upwind spatial discretization. Solution convergence was monitored by minimizing the residuals for the mass continuity and velocity components. For pressure gradient-driven laminar flow solutions, the typical execution time was 1–2 days for 20,000 iterations with 0.0001 sec time-step sizes using parallel computations with four processors operating at 3.1 GHz on Xeon nodes. The pressure and velocity data at the point (around the fenestra opening) adjacent to the blood vessel depression were collected from this solution in order to use them as input field parameters in the subsequent simulation of plasma perfusion inside the tumor region. The density and viscosity of the plasma (Nithiarasu 2022) were set at 1030 kg/m^3^ and 0.001 kg/m.s, respectively. The density and viscosity of WBCs were determined to be 1080 kg/m^3^ and 0.011 kg/m.s, respectively. On the other hand, the viscosity of RBCs is determined by our assigned functions, and the density of RBCs is set at 1100 kg/m^3^. The shape factor influences the phase interaction between plasma and RBCs (through our custom-made functions). The diameters of RBCs and WBCs (Schwartz, R. S. and Conley, C. L. 2020) are set to 7 *μ*m and 14 *μ*m, respectively.

### 2.3. Simulation of plasma perfusion inside the tumor region

The mechanics of plasma transport through the fenestra are tracked as a viscouslaminar-transient state flow. We obtained the numerical solution by employing the Eulerian multiphase model, which eliminates the need for custom interparticle phenomena equations. Herein, plasma is the primary phase, whereas air is the secondary phase.

The numerical solution’s convergence was again determined by minimizing the residuals of the mass and velocity components. For pressure gradient-driven laminar flow solutions, typical execution time 3–4 hours for 3000 iterations with 0.0001 sec time-steps (for Model 1, Model 2, and Model 3), using 4-processor-based parallel computations operating at 3.1 GHz on Xeon nodes. Amongst additional physical parameters, the density and viscosity for air is assumed to be 1.225 kg/m^3^ and 1.7894 × 10^−5^ kg/m.s, respectively. The diameter of air particle is set to 4.12 × 10^−10^ m (Porterfield, W. W. and Kruse, W. 1995). Note that our group has previously employed similar validated computational tools to replicate air and particle transport in respiratory physiology (e.g., Basu, S. 2021, Basu, S. and Frank-Ito, D. O.and Kimbell, J. S. 2018, Basu, S. et al. 2020).

### 2.4. Boundary conditions enforced

During the simulation of the blood vessel region (see Section 2 2.2), the inlet pressure (Zhao, G. et al. 2007) is set to 3325 Pa and is designated as the pressure inlet, the outlet pressure (Zhao, G. et al. 2007) is set to 2128 Pa and is designated as the pressure outlet, and the streamwise vessel extents have no-slip boundary condition. Further, at the blood vessel outlet, the back flow volume fraction is set to 0 for WBCs and 1 for RBCs, respectively. During the simulation of the tumor region (see Section 2 2.3), the inlet pressure for the mixed phases is set to the pressure values obtained from the blood vessel simulation, and the velocity is also set to the values obtained from the blood vessel simulation via user-defined commands. As a boundary condition, the air outlet pressure is set at 2780 Pa which is extrapolated from an earlier study (Wu, M. et al. 2013) and our extrapolated value agrees with these high interstitial pressure findings (Heldin, C. et al. 2004, Stylianopoulos et al. 2018, Sven, K. and Josipa, F. 2007) inside tumor region and is designated as a pressure outlet.

## 3. Results

### 3.1. Simulated transport parameters inside the blood vessel

Analysis of the volume fraction trends from the blood vessel simulation reveals that plasma has a constituent volume fraction greater than 0.50 and less than 0.60, RBCs have a constituent volume fraction greater than 0.40 and less than 0.50, and WBCs have a constituent volume fraction less than 0.10. Figure 4 depicts these scenarios, which are the same for different flow time intervals of 0.10, 0.15, and 0.30 sec. These findings are consistent with earlier findings (McWhirter, J.L. and and Gompper,G. 2012). Also, the resulting plasma pressure and velocity distribution at the fenestra opening provided the inputs for simulating intra-tumoral plasma uptake, as discussed next.

**Fig. 4:**
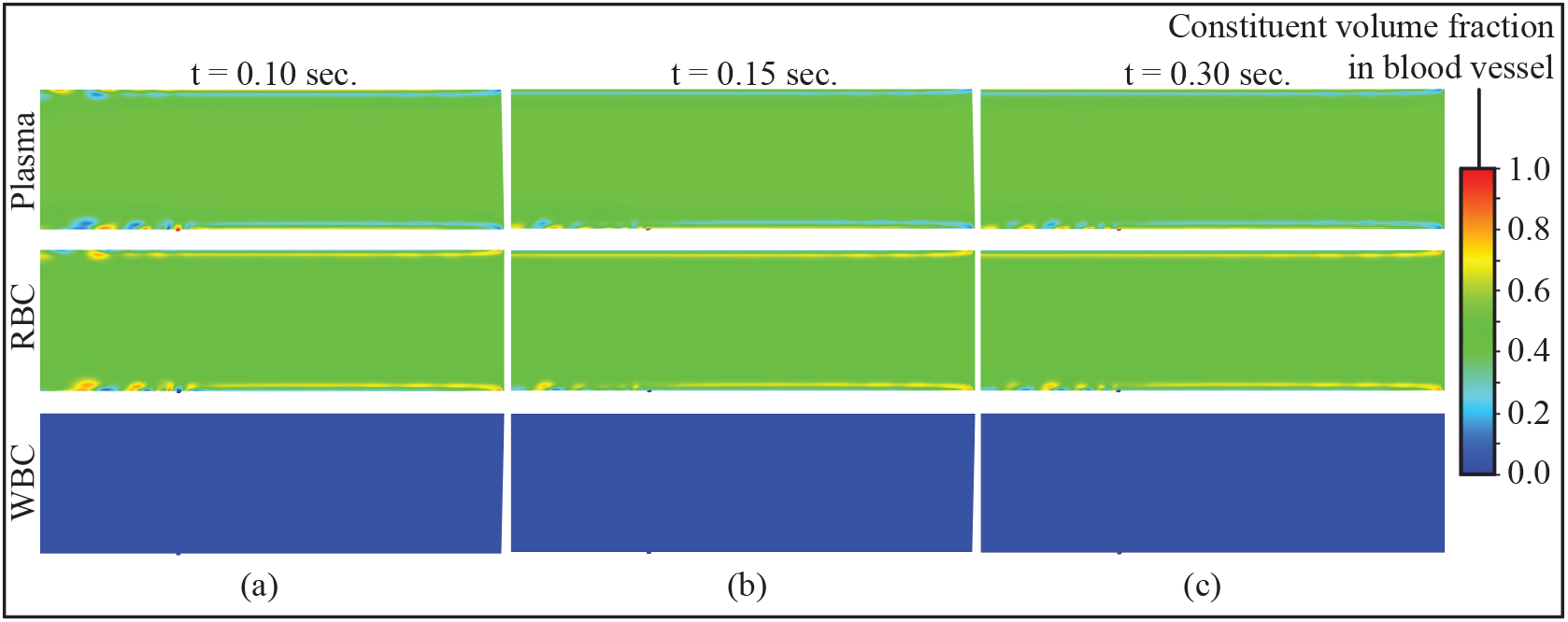
Panels (a)-(c) show the blood flow scenario with 3 phases inside the blood vessel with depression; three phases are: Plasma, RBCs (Red Blood Cells), WBCs (White Blood Cells). Panel (a) shows the state of 3 phases when flow time is 0.10 sec, and panels (b)-(c) respectively show states at 0.15 sec and 0.30 sec. Panels (a)-(c) depict the volume fractions of Plasma, RBCs, and WBCs in blood vessel. Color scheme: Deep green zone volume fraction range: 0.50 − 0.58, Light green zone volume fraction range : 0.35 − 0.49, Dark blue zone volume fraction range : 0.00 − 0.10.

### 3.2. Perfusion of plasma into the tumor vasculature

Eulerian 2-phase simulations with plasma as the primary phase and air as the secondary phase quantified the perfusion trends in the tumor extracellular space. The pressure and velocity measurements obtained at each time step of the simulations in the main blood vessel are used as transient inlet boundary conditions for the tumor region. When the diameter of the fenestra remains constant, we can observe the following: in the first row (see Figure 5), the fenestra diameter is 0.1 *μ*m, but the flow time increases from left to right, resulting in increased plasma percolation (indicated by the red zones in Figure 5). When we examine the first column of Figure 5, we notice a similar trend of plasma percolation, where the diameter of the fenestra varies from 0.1 to 0.3 and then to 0.5 *μ*m, but the flow duration of 0.10 sec remains constant. Thus, leakiness of the plasma increases progressively as the fenestra dimension and flow time increase. We can observe that at 0.30 sec flow time (Model 2, Model 3), the ratio of plasma to air is larger than one (see Table 1), indicating that plasma has approached its percolation limit and has mostly replaced air in the tumor.

**Fig. 5:**
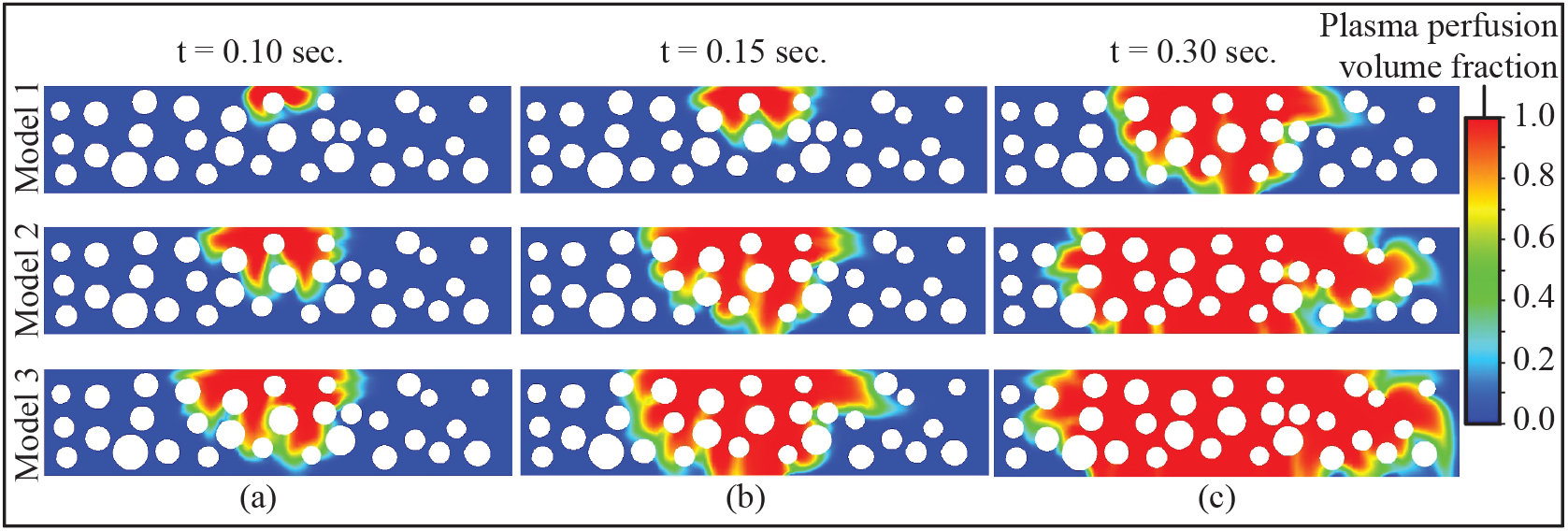
Panels (a)–(c) illustrate the plasma leakiness trend for three distinct flow times into the tumor domain. Model 1 has a fenestra size of 0.1 *μ*m, Model 2 has a fenestra size of 0.3 *μ*m, and the Model 3 has a fenestra size of 0.5 *μ*m. The blue zoned area is dominated by air, while the red and green zoned areas denote the plasma leakiness trend within the tumor region.

**TABLE 1:** Color palette analysis of Figure 5: A spatio-temporal representation of plasma perfusion trend into the tumor extracellular space – represented by the ratio of the summation of red (R) and green (G) domains over the blue (B) region.

|  | Flow time = 0.10 sec | Flow time = 0.15 sec | Flow time = 0.30 sec |
| --- | --- | --- | --- |
| Model 1 | (R+G):B = 0.021 : 1 | (R+G):B = 0.024 : 1 | (R+G):B = 0.42 : 1 |
| Model 2 | (R+G):B = 0.087 : 1 | (R+G):B = 0.167 : 1 | (R+G):B = 4.10 : 1 |
| Model 3 | (R+G):B = 0.141 : 1 | (R+G):B = 0.219 : 1 | (R+G):B = 4.65 : 1 |

## 4. Discussion

- *On the limitations of biomimetic modeling –* The diameter and depth of the depression were chosen such that it remains wider (i.e., its diameter is 5.2 *μ*m) than the maximum diameter of the fenestra (0.5 *μ*m), implying that the fenestra should lie within the depression. Additionally, the depression’s depth is 2.6 *μ*m, which is greater than the fenestra’s height of 0.3 *μ*m. However, these values were chosen arbitrarily in order to simplify the biomimetic model (see Section 2 2.1). The biomimetic model is divided into two segments for the primary reason of algorithmically restricting the entry of larger particles (the particle diameter being greater than the diameter of the fenestra) into the tumor via the fenestra. Also, the numerical scheme has employed user-defined functions built based on earlier published studies Jung, J. and Hassanein, A. 2008, and Jung, J. et al. 2006, and has been compared for validation against the experimental results from Karino, T. and Goldsmith, H. L. 1977.
- *On the limitations of simulating the plasma perfusion inside the tumor region –* In our reduced-order modeling, we did not consider plasma transportation within the enitre volume of the tumor; rather, the focus was on a small chunk of tumor in order to quantify the leakiness trend of plasma within the tumor region. The outcome is summarized in Table 1.
- *On the relation between the fensetra diameter and the rate of diffusion –* By analyzing the spatio-temporal plasma perfusion trends, represented through the color palette data in Table 1, we can infer that for the same flow time, the combined area of red and green zones (see Figure 5) increases relative to the blue zone area as the diameter increases (see Section 3 3.2). The combined area of red and green represents plasma percolation, while the blue represents air. In the following, we show how the size of the opening of the fenestra affects the rate at which plasma moves through the tumor.

At flow time j sec,

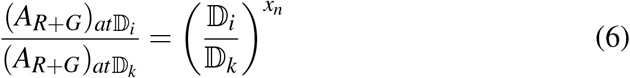

Here, *A*_*R*+*G*_ represents the combined area of red and green (see Table 1) and D indicates the diameter of fenestra, *i* and *k* correspond to the different sizes of diameters, j indicates the flow time, and *x*_*n*_ indicates the power of the fenestra diameter ratio for specific flow time.

Therefore, as a representative case, for flow time = 0.1 sec:

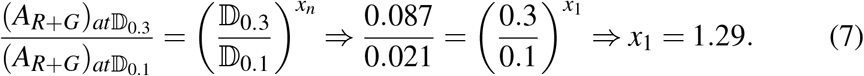

Table 2 depicts the power estimates (derived following the same strategy as above) for different flow times (e.g., 0.15 sec, 0.30 sec) and diameter ratios. The slope is 4 between (0.10, 1.09) and (0.15, 1.29), and 0.67 between (0.30, 1.39) and (0.10, 1.29). As can be seen, the slope is decreasing, indicating that the rate of diffusion is decreasing in lockstep with the increase in diffusion distance as flow time increases. The plasma percolates away from the fenestra opening inside the tumor as the flow increases, implying that the distance between them grows. This scenario lends credence to Fick’s law. According to Fick’s law (Bouzin, C. and Feron, O. 2007, West, J. B. 2012), the rate of diffusion is inversely proportional to the distance over which it occurs.

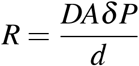

Here, *R* = rate of diffusion, *d* = distance over which diffusion occurs, *A* = area over which diffusion occurs, *δ P* indicates the pressure difference, *D* = diffusion coefficient. So, *R* ∝ 1/*d*; which implies that rate of diffusion will decrease if the perfusion distance increases.

**TABLE 2:** Ratio of the power for the different fenestra diameter ratios to estimate perfusion trend comparisons over different flow times.

| Flow time (sec) | the power ($x$) for $\frac{\mathbb{D}_{0.3}}{\mathbb{D}_{0.1}}$ | the power ($y$) for $\frac{\mathbb{D}_{0.5}}{\mathbb{D}_{0.1}}$ | $\frac{x}{y}$ |
| --- | --- | --- | --- |
| 0.10 | 1.29 | 1.18 | 1.09 |
| 0.15 | 1.77 | 1.37 | 1.29 |
| 0.30 | 2.07 | 1.49 | 1.39 |

- *On inputs to the CT (Computed Tomography) scan of mice –* Human pancreatic cancer cells were orthotopically injected into the pancreas of nude mice to establish human pancreatic tumors. To assess a realistic scenario wherein our modeling strategies can find applicability, pancreatic orthotopic tissues from 3 nude mice were collected and fixed for 24 hours in formaldehyde. Paraffin-embedded 5 *μ*m thick sections of tumor tissues were prepared. Sections were deparaffinized with Histo-Clear and ethanol. Then tissue slides were rehydrated and stained with hematoxylin for 5 min. The tissue slides were then washed with distilled water, soaked in 95% ethanol for 30 sec, and stained with eosin for 1 min. Next, they were dehydrated with 100% ethanol for 1 min, washed in xylene and mounted with a coverslip using a Hardset Mounting Medium. Finally, slides were visualized using a Zeiss inverted Axio Observer Z1 microscope.

To determine whether our STL (Stereolithography) imaging reconstruction can accurately replicate the overall architecture of tumor tissue (thus easing the first step of the computational model generation process), we compared a stained human pancreatic orthotopic cancer tissue sample (with hematoxylin and eosin, or H&E staining) to the STL version of the same section of the stained cancer tissue sample. The H&E stain technique, the most commonly used modality in medical diagnosis, especially by pathologists for cancer diagnosis, provides information on overall cell structure, patterns, and shapes in a tissue sample. We observe that the overall architecture as well as the microanatomy of the stained tumor tissue section (see panel (a) in Figure 6) is nearly identical to the tissue scan from the STL image (panel (b) in Figure 6). For example, the necrotic area in both images (see Figure 6) were nearly identical in size and pattern. Furthermore, the STL geometry accurately pinpointed the cells undergoing apoptosis / swollen cells (see Figure 6). We also noted that the STL reconstruction recognized the cellular nuclei within the tumor samples. Collectively, this preliminary assessment demonstrated that our explained *in silico* can accurately replicate the overall architecture of human pancreatic orthotopic tissue samples.

**Fig. 6:**
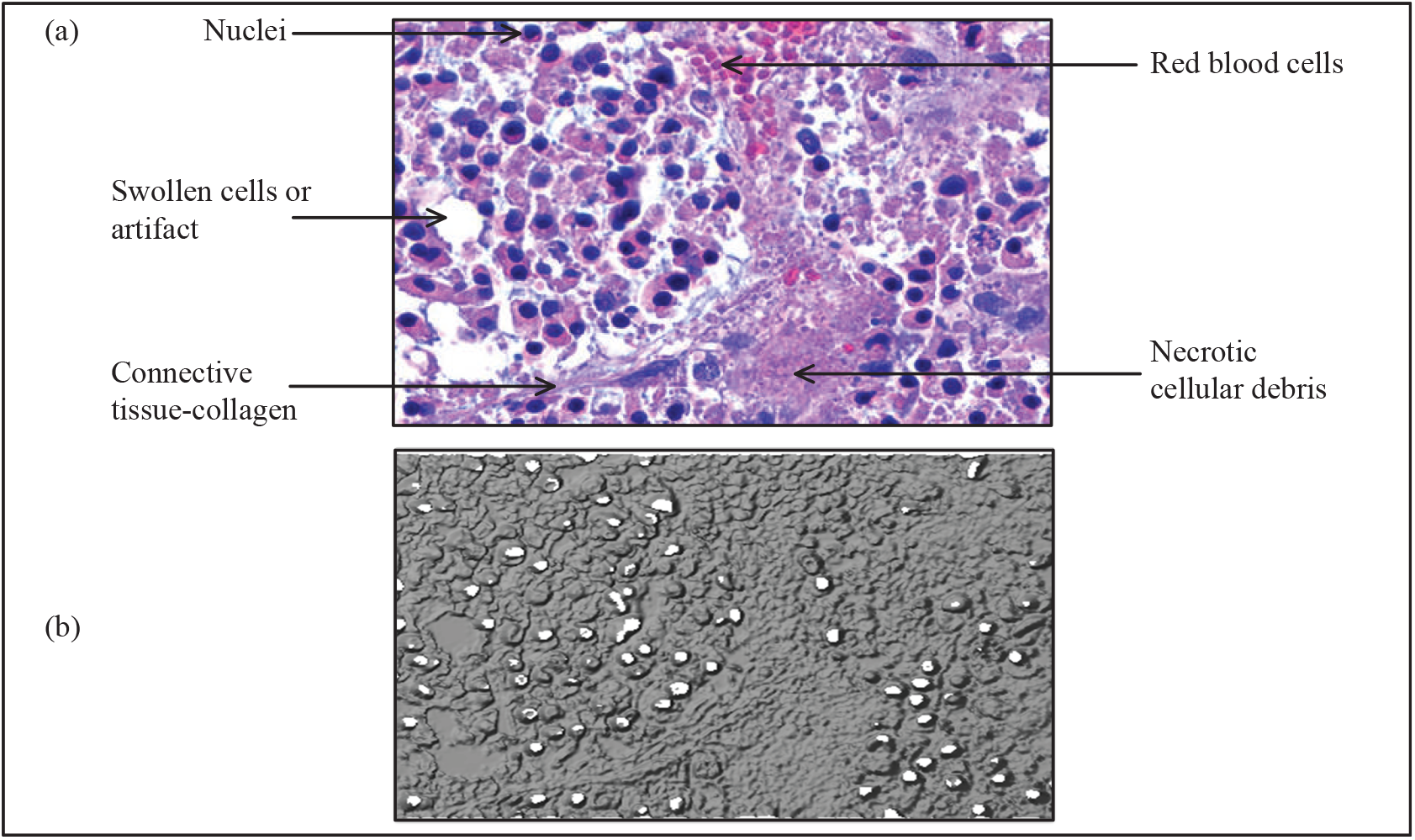
Panel (a) is the representative image from H&E (hematoxylin and eosin) staining done to identify overall architecture (shape, pattern, and structure of cells) of human pancreatic orthotopic tissue. The staining was performed on pancreatic tissue from three different samples. Panel (b) shows the STL (Stereolithography) reconstruction of panel (a).

## 5. Conclusion

Diffusive transport into solid tumors is unlikely due to their dense fibrous extracellular structure, whereas convection is difficult owing to a lack of blood and lymphatic movements. The complexity of fluidic transport inside the tumor microenvironment comes with unanswered questions with potentially significant translational implications. For this study, we have built a 2D biomimetic model of a blood vessel with an adhering reduced-order tumor extracellular domain with fenestra openings to the vasculature. In our CFD-based work, plasma percolation is found to be proportional to the leakiness of the fenestra aperture and to the increase in flow duration tracked. Although we conducted our study on a small chunk of the tumor, the study’s outcome indicates that it can be applied to the entire tumor region. In future, this CFD transportation model can be applied to the real tumor vasculature; for this purpose, we have already stained real mice tumors (see Section 4) and constructed the STL of that tumor (see Figure 6).

In summary, the derived plasma penetration levels are clearly impacted by tumor leakiness (modulated by varying the fenestra openings). This work can have key implications for *in silico* diagnosis of hard-to-reach tumor types using computational models built from medical imaging data.

## Acknowledgements

The authors thank the *Experimental and Computational Multiphase Flow* editorial team for the invitation to publish in the special issue on “Multiphase flow application in the human body”.

## Author contributions

MMHA: geometry generation, numerical simulations, data curation, investigation, writing; NC: geometry generation, numerical simulations, data curation; JM: tumor scanning, writing; KR: tumor scanning, student supervision; SB: conceptualization, funding acquisition, project administration, student supervision, writing.

## Data availability

The reduced biomimetic geometries, the simulation data sets – are available onrequest from the corresponding author, through a shared-acces <u>Google Drive link</u>.

## Funding note

The reported work was supported by a National Institutes of Health (NIH) CO-BRE Pilot Grant from the North Dakota State University Center for Diagnostic and Therapeutic Strategies in Pancreatic Cancer – Project Number 5P20GM109024. Any opinions, findings, and conclusions or recommendations expressed here are, however, those of the authors and do not necessarily reflect views of the NIH.

